# Replicating Arabidopsis Model Leaf Surfaces for Phyllosphere Microbiology

**DOI:** 10.1101/523985

**Authors:** Rebecca Soffe, Michal Bernach, Mitja Remus-Emsermann, Volker Nock

**Affiliations:** Department of Electrical and Computer Engineering, University of Canterbury, Christchurch, New Zealand; School of Biological Sciences, University of Canterbury, Christchurch, New Zealand

**Keywords:** Microfabrication, PDMS, Biomimetic Surface, Replica Molding, Phyllosphere, Microbiology, Arabidopsis

## Abstract

Artificial surfaces are commonly used in place of leaves in phyllosphere microbiology to study microbial behaviour on plant leaf surfaces. Studies looking into individual environmental factors influencing microorganisms are routinely carried out using artificial surfaces. Commonly used artificial surfaces include nutrient agar, isolated leaf cuticles, and reconstituted leaf waxes. However, interest is growing in using microstructured surfaces mimicking the complex topography of leaf surfaces for phyllosphere microbiology. As such replica leaf surfaces, produced by microfabrication, are appearing in literature. Replica leaf surfaces have been produced in agar, epoxy, polystyrene, and polydimethylsiloxane (PDMS). However, these protocols are not suitable for replicating fragile leaves such as of the model plant *Arabidopsis thaliana*. This is of importance as *A. thaliana* is a model system for molecular plant genetics, molecular plant biology, and microbial ecology. Here we present a versatile replication protocol for replicating fragile leaf surfaces into PDMS. We display the capacity of our replication process using optical microscopy, atomic force microscopy (AFM), and contact angle measurements to compare living and PDMS replica *A. thaliana* leaf surfaces. To highlight the use of our replica leaf surfaces for phyllosphere microbiology, we visualised bacteria on the replica leaf surfaces in comparison to living leaf surfaces.

## Introduction

Researchers often turn to nature for inspiration in developing new and innovative technologies.^1^ For instance, taking inspiration from the colour changing ability of chameleons, to hummingbird and dragonfly inspired micro/unmanned air vehicles, or replicating surfaces for their different physical properties.^2-5^ In addition, researchers also mimic nature, to provide more insight into our natural world. For instance, creating organs-on-a-chip for *in vitro* studies to minimise the use of animal surrogates, to developing microfluidic platforms for physiological studies, or developing artificial surfaces for phyllosphere microbiology.^6-14^

In phyllosphere microbiology, the study of microorganisms that reside on the leaves of plants, artificial surfaces are used to provide an insight into microbial behaviour in the phyllosphere. The use of artificial surfaces in place of a living leaf, enable a reductionist approach, to investigate the impact of individual factors on microorganism functioning and viability, for example.^14^ Artificial surfaces include: (1) flat surfaces, such as nutrient agar, and inert surfaces (for example stainless steel); and (2) microstructured surfaces, such as isolated leaf cuticles, leaf peels, reconstituted leaf waxes, and microfabricated surfaces.^14-20^ Although these surfaces are suitable for their intended purpose, they do not suitability represent the complex topography of plant leave required for some microorganism studies.^21,22^ Thus, protocols utilizing double-casting approaches, have been reported to develop leaf replicas in agar, epoxy, polystyrene, and polydimethylsiloxane (PDMS).^22-26^

Fabricating replica surfaces inspired by nature, such as leaves have primarily been focused on replicating their superhydrophobic, superoleophobic, and superhydrophilic properties.^2-5^ Such surfaces have applications in anti-reflection, anti-fouling, and anti-fogging coatings.^5,26,27^ For example, Schulte *et al.* replicated leaves for new biomimetic surfaces with new optical properties.^24^ However, recently replicating leaf surfaces for phyllosphere microbiology has gathered interest. This is attributed to an increase in interest in the role that microorganisms have on plant health, for example, foliar diseases.^14^ In addition, unwanted microorganisms in the phyllosphere can arise due to the exposure of potential contamination sources during plant growth on a farm. This is important for leafy greens such as spinach, rhubarb, and parsley that is produced for human consumption. In addition, they are consumed either raw or without minimal processing; thus, unwanted contamination is neither removed nor killed.^*11-14,28-31*^ On the rare occasion, contamination can lead to outbreaks that can result in severe illnesses.^32-34^ Hence, new studies are imperative for increased understanding of phyllosphere microbiology, which will enable mitigation strategies to be developed.^34,35^

In one study, leaf replicas in epoxy were used to investigate the influence of cuticular folds on the behaviour of Colorado potato beetles (*Leptinotarsa decemlineata*).^36^ In another study, Zhang *et al.* replicated spinach leaf surfaces, to produce replicas in agarose and PDMS. They investigated the influence of microstructures on the behaviour of *Escherichia coli* on both flat and replica spinach leaf surfaces made from agarose.^23^ Furthermore, we have recently reported that leaf replicas produced in PDMS are more suitable for phyllosphere microbiology compared to those produced in agarose and gelatin. We came to this conclusion by investigating the resolution, degradation, contact angles, and bacterial survival on the three aforementioned materials. Such that, the replica produced in PDMS were of a high fidelity and did not degrade under different humidity conditions, had comparable hydrophobicity to two generic isolated leaf cuticles, and bacteria did not survive which was in agreeance to the observations made on the isolated leaf cuticles.^15^

For our model leaf, we decided to work with *Arabidopsis thaliana*, which is the best-established model system for molecular plant genetics, and molecular plant biology. In addition, *A. thaliana* is a well-established model system for microbial ecology.^11,37,38^ The downside of *A. thaliana* is that the leaves are inherently fragile, which makes the leaves difficult to replicate. In addition, the plants we used were grown in optimal conditions in either soil or media culture in growth chambers. In contrast, reported protocols have been using plants grown in the wild (for example in a botanical garden),^24^ in less optimal conditions, such as inconsistent water supply and uncontrolled temperature and light conditions. Growing plants in optimal conditions, especially under tissue culture conditions results in: (1) the leaves holding a higher amount of moisture; and (2) a reduced amount of leaf cuticular wax.^39,40^ Thus, providing a challenge to imprint the leaf surface with high fidelity. For example, imprinting a leaf with polyvinylsiloxane does not allow for degassing to maximise resolution, due to the quick curing time.^24,36^ Light curable polymers have also been utilised, which results in undesirable chemical and ultraviolet light exposure to the plant leaf.^41^ In addition, protocols have also been reported using PDMS to imprint leaf surfaces and produce leaf replicas in PDMS with good resolution.^23,26,42^ However, these protocols have some drawbacks: (1) the leaf can be exposed to excessive heat during curing, which results in the leaf shrivelling during curing, and subsequent loss of fidelity of the replica; (2) the leaf samples retain too much water, which results in the PDMS not curing on the leaf surface; and (3) leaf residue can remain affixed to the imprint.^23,26,42,43^ This in turn affects the quality of the imprint and subsequent replica.^41^

As a result, we developed a versatile protocol to maximise the fidelity of the leaf imprint, and subsequent replica leaf. For the PDMS imprint, we use a base to curing agent ratio of 20:1. Whereas, published procedures for producing leaf replicas with PDMS, use a PDMS ratio of 10:1 for both the imprint and replica.^23,42^ A ratio of 10:1 is also commonly used in other double-casting applications, such as bioimprinting.^44-46^ Prior to casting the leaf imprint the *A. thaliana* samples were briefly dried to remove the surface moisture of the leaves. This was done to ensure the PDMS cures on the surface. Furthermore, the PDMS imprint was cured at 45 °C for 20 h. Due to the concern of leaf residue affecting the resolution of the replica, the PDMS imprint was placed in a leaf digestion solution to remove any leaf residue. An anti-adhesion layer was then applied to the imprint, to enable a PDMS replica leaf to be peeled off the imprint. The replica leaf was produced at a ratio of 10:1.

We highlight the capacity of the replication process using optical microscopy, atomic force microscopy (AFM), and contact angle measurements. With replica and living leaf surfaces of *A. thaliana* compared. To highlight the use of our replica procedure to produce replica leaf surfaces for phyllosphere microbiology, we introduced bacteria onto their surfaces

## Methods and Materials

### Growth Protocol for *A. thaliana* Plants Grown in Soil

Plastic plant pots (70 by 70 by 90 mm) were filled with potting mix containing 9 month-controlled release fertiliser (Canterbury Landscape, New Zealand), 20 mm below the top of the pots. A 2 g scoop of No Insects Lawngard Prills (KiwiCare, New Zealand) was added to the soil. The last 20 mm of the plant pots were filled with potting mix sieved with 5 mm mesh. The *A. thaliana* ecotype Columbia 0 (Col-0) wild type seeds were suspended in water, and then placed on top of the the soil in the plant pot. The plant pots were then placed on a raised tray with holes, to allow the water to be gradually be absorbed by the soil in the plant pots. This tray was then placed in a container with a clear cover containing holes. The plants were grown under long day conditions as follows: 16 h of daylight at 21 °C and 8 h of darkness at 18 °C, and the plants were watered weekly from initial potting. Seedlings were trimmed to ensure that only one seedling remained. Furthermore, the cover of the container was removed after three weeks.

Two days prior to replication of the leaves surfaces, the plants were removed from the growth chamber and placed on the bench in a climate-controlled laboratory. This was done to reduce the internal water content of the plants. This was undertaken to reduce the amount of moisture being released from the leaf samples, while the PDMS was curing.

### Growth Protocol for *A. thaliana* Plants Grown on Culture Media

Initially the *A. thaliana* seeds were sterilised using a standard protocol, as follows: (1) In a 1.5 mL Eppendorf tube the desired amount of *A. thaliana* Columbia (Col-0) wild type seeds were added; (2) Then 1 mL of 50% bleach (Dynawhite, Jasol, New Zealand) was added to the Eppendorf tube; (3) After five minutes, the bleach was removed from Eppendorf tube; (4) Then 1 mL of 70% ethanol (Anchor) was added to the Eppendorf tube; (5) After one minute the ethanol was removed, and the seeds were washed five times with 1 mL of sterile deionised water.^47^ After sterilisation, the seeds were stratified at 4 °C in sterile deionised water for at least two days in the dark.

Glass jars (741 Mold Jar, Weck Jars, Germany) were used to grow the *A. thaliana* plants on culture media. Prior to adding the seeds and culture media, the glass gars were sterilised by autoclaving. Then 65 mL of Murashige and Skoog medium (pH 5.8, M0222, Duchefa Biochemie, Netherlands) with 0.6 % w/v plant agar (P1001, Duchefa Biochemie, Netherlands), and 1 % w/v of Sucrose (Product info, company) were added to the jars. The open jars were then placed in laminar flow for an hour, to cure the agar and dry the medium. In addition, to further minimise the potential occurrence of contamination, the media filled weck jars were ultraviolet sterilised for 20 minutes under laminar flow. The *A. thaliana* seeds were then placed on the surface of the culture media. The jars were then placed inside a growth chamber (Contherm Precision Environmental Chamber, Contherm Scientific, New Zealand). A long day growth condition was used: 16 h of daylight at 21 °C and 8 h of darkness at 18 °C.

### Preparation of *A. thaliana* Leaf Samples

Double-sided mounting tape (Orabond 1397PP, ORAFOL Europe GmbH, Germany), was placed on to the bottom of a polystyrene petri dish. Care was taken to ensure that the tape was sufficiently flatten, to minimise affecting the quality of the imprint and subsequent replica. In addition, the tape should be larger than the size of the leaf. Using two pieces of tape side by side caused uneven surfaces from the placement of the tape, and bubbles forming in the gap between the two pieces of tape. Leaves were taken from the *A. thaliana* plants, rinsed with deionised water to remove any contaminants, and promptly dried with low pressure dry nitrogen. The leaves were either kept whole, or cut into smaller samples, and placed on the double-sided tape. The leaves were then carefully flattened by pressing gently on the leaves with softened Parafilm “M” (Bemis, USA) (**Figure 1a**). The samples were then placed in an automatic desiccator (Secador 2.0, SP Scienceware Bel-Art, USA) or a vacuum desiccator (Z119016, Sigma-Aldrich) until no residual moisture was observed on the surface of the leaf samples. This process was selected to minimise damage to the microstructures of the leaf samples, and remove any excess moisture from the top surfaces of the leaves that would prevent the PDMS from setting.^48^

**Figure 1:**
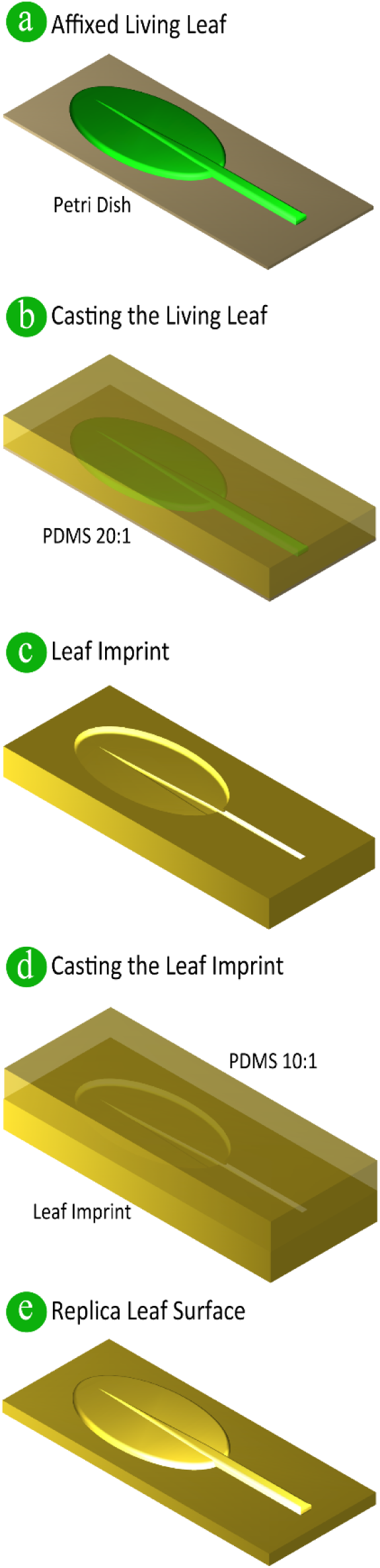
Schematic of the Replication Protocol for Replica Leaf Surfaces in PDMS. (a) Affix a living leaf sample to a petri dish with double-sided tape. (b) PDMS with a ratio of 10:1 is then poured on to the affixed leaf, with the PDMS degassed and cured. (c) The leaf imprint is then carefully peeled off. Prior to pouring the leaf imprint with PDMS, an anti-adhesion coating is applied. (d) PDMS with a ratio of 10:1 is poured onto the imprint, with the PDMS being degassed and cured. (e) Once cured the leaf replica is peeled carefully from the leaf imprint.

### Leaf Imprint Protocol

The polydimethylsiloxane (PDMS) (Sylgard 184, Dow Corning) was prepared at a ratio of 20:1 w/w, base to curing agent.^43^ The PDMS was thoroughly mixed, then degassed for 20 to 30 minutes in a vacuum desiccator (Z119016, Sigma-Aldrich), until no bubbles remained. The PDMS was then promptly poured onto the leaves affixed to the petri dish (**Figure 1b**). The PDMS was degassed for an hour, or slightly longer if any bubbles remained. The petri dish then was placed in an oven at 45 °C for 20 h – to cure the PDMS. Curing for a longer duration resulted in an increase in leaf residue remaining on the imprint. We conjecture this was due to the degradation of the leaf cuticle.^43^ Once the PDMS was cured the petri dish was removed from the oven, and left to cool to room temperature. Once cool, the imprint was carefully peeled from the leaf sample (**Figure 1c**).

### Removing Leaf Residue from the PDMS Imprint

To remove any leaf residue, a digestion solution comprising of 3.5% w/v sodium hydroxide (NaOH, S5881, Sigma-Aldrich) and 2.5% w/v sodium carbonate (Na_2_CO_3_, 222321, Sigma-Aldrich), in deionised water was used.^49^ The digestion solution was placed on a hotplate at 160 °C and stirred at low angular velocity (80 to 120 rpm, depending on the size of the glassware and magnetic stirrer used) (SP88857105, Cimarec+, Thermo Scientific), until the sodium hydroxide and sodium carbonate was completely dissolved – approximately 15 minutes. The PDMS leaf imprint was placed into the digestion solution for 20 minutes. Once removed from the digestion solution, the PDMS leaf imprint was promptly rinsed thoroughly with deionised water. Any stubborn residue was carefully removed with tweezers with the PDMS leaf imprint in deionised water (usually this occurs around the edges of the leaf sample). The PDMS leaf imprint was then placed in deionised water for 20 minutes. The imprint was then rinsed thoroughly with deionised water and dried with dry nitrogen.

### Anti-adhesion Coating

An anti-adhesion coating can be produced in one of two ways: (1) Treatment with hydroxypropylmethylcellulose (HPMC, H8384, Sigma-Aldrich). The leaf imprint was treated with 0.3% w/v HPMC in phosphate buffer saline (PBS, P4417, Sigma-Aldrich) for 10 minutes for anti-adhesion. The imprint was then rinsed with deionised water, and subsequently dried with nitrogen.^45,46^ Alternatively, (2) treatment with trichloro(1H,1H,2H,2H-perfluorooctyl)silane (448931, Sigma-Aldrich), can be undertaken. The imprint was initially placed in oxygen plasma for 60 s to produce a hydrophilic surface (PIE Scientific Tergeo Plasma Cleaner, USA). The imprint was then placed in a vacuum desiccator (Z119016, Sigma-Aldrich) alongside a bottle containing a small droplet of trichloro(1H,1H,2H,2H-perfluorooctyl)silane and placed under vacuum for an hour.^50,51^

### Replica Leaf in PDMS Protocol

For the PDMS replica leaf, the base and curing agent were thoroughly mixed together, at a ratio of 10:1 w/w (base to curing agent). The PDMS was then degassed in a vacuum desiccator (Z119016, Sigma-Aldrich) until no bubbles remained. The PDMS was subsequently poured onto the anti-adhesion coated leaf imprint and degassed for an hour (**Figure 1d**). The PDMS was placed on a hotplate for two hours at 80 °C to cure. Finally, the PDMS replica leaf was carefully peeled off the leaf imprint, once the PDMS cooled to room temperature (**Figure 1e**).

### Atomic Force Microscopy (AFM) Imaging

All AFM images were taken using a MFP-3D Origin (Asylum Research - Oxford Instruments, USA), equipped with either TAP150Al-G (*A. thaliana*) or TAP300-G (PDMS imprint and replica) tips (BudgetSensors, USA) operating in tapping mode. All AFM scans were analysed using Gwyddion (Version 2.49).

### Contact Angle Measurements

Contact angle measurements were obtained using a CAM200 (KSV Instruments Ltd, Finland), integrated with KSV CAM Optical Contact Angle and Pendant Drop Surface Tension Software (ver. 4.01, KSV Instruments Ltd, Finland). Deionized water was used to determine the surface energy of the *A. thaliana* and PDMS leaf replica. Five samples were measured for both *A. thaliana* and PDMS replica leaves. In all cases the samples were obtained using a cork borer (Usbeck, Germany) with a diameter of 11.5 mm. Water droplets with a volume less than 60 µl were recorded.

Results are presented as mean ± SEM (standard error of the mean). For statistical analysis, two-way ANOVA was performed using GraphPad Prism 7 (GraphPad Software, USA). P values less than 0.05 were considered significant (*P<0.05, **P<0.01, ***P<0.001, and ****P<0.0001).

### Bacteria *Pantoea agglomerans* 299R∷MRE-Tn7-145 Culture Protocol

*P. agglomerans* 299R∷MRE-Tn7-145, a leaf colonising bacterium was grown overnight on nutrient agar plates (13 gL^-1^ Lysogeny broth and 15 gL^-1^ bacteriological Agar, Oxoid) containing 20 mg/L of gentamycin (AG Scientific) at 30 °C.^52^ The bacteria was then harvested using a sterile inoculation loop and was suspended in 5 ml of sterile phosphate buffer saline (PBS pH 7.4, P4417, Sigma-Aldrich). The bacteria was then washed by centrifugation at 1150 RCF for five minutes at 10 °C. The supernatant was discarded, and the bacteria was suspended in fresh phosphate buffer, and washed for a second time – following the aforementioned process. After washing for a second time the bacteria were suspended in PBS to an OD_600 nm_ of 0.7.

### Bacteria Visualisation Protocol

Living *A. thaliana* leaves and PDMS replica leaves require different sample preparation for bacteria visualisation studies, undertaken using microscopy.

The *A. thaliana* leaf samples were taken from mature *A. thaliana plants,* immediately prior to inoculation of bacteria and subsequent microscopy. The abaxial leaf samples were first washed with deionised water to remove any contaminants. The leaf samples were then dried with either filtered compressed air or nitrogen. For microscopy the leaf samples were affixed to glass microscopy slides with double-sided tape.

Whereas, the PDMS replica leaves were sterilised for 15 minutes using ultra-violet sterilisation. Once sterilised the replica leaves were placed in vacuum desiccator (Z119016, Sigma-Aldrich) for two hours. The replicas were then placed in PBS overnight at room temperature. Prior to the inoculation with bacteria the PDMS replicas were dried with either filtered compressed air or nitrogen.

Prior to microscopy two aliquots of 200 µl of bacterial solution were inoculated onto the living *A. thaliana* or PDMS leaf replica samples using an airbrush (KKmoon T-180 Airbrush, China) at 1×10^5^ Pa.^53^ Differential interference contrast (DIC) and fluorescent images were then obtained using a Zeiss Axiolmager M1 fluorescent widefield microscope equipped with a 43HE Zeiss filter set (Zeiss, Germany). Images were acquired using a 20 × objective (Zeiss, Germany), and an Axiocam 506 camera (Zeiss, Germany) controlled by Zeiss Zen software (version 2.3). The DIC and fluorescent images were obtained using DIC 2 (1.5 ms exposure time) and the red fluorescent channel (850 ms exposure time), respectively. Images were then processed using Fiji (version 1.52 h), and the extended depth of focus plug-in was utilised to process the DIC channel.^54^

## Results and Discussion

### Replication Protocol

The replication protocol displayed in **Figure 1** was used to replicate the leaves of *Arabidopsis thaliana* plants, which is a well-established model system for microbial ecology.^37^ For our replication protocol we used leaf samples from mature *A. thaliana* plants that had been grown for four to six weeks. We successfully replicated the surface topography of plants grown in either soil or media culture. The ability to replicate plants grown in either condition is important, as the growth conditions are dependent on the studies that will be undertaken. For studies focusing on the plant, the plants will be grown in soil. However, if the plants are required to be axenic (free from living microorganism) they are grown on sterile culture media, which is used for investigating microorganism communities.^55^

Initially, to produce the polydimethylsiloxane (PDMS) leaf imprints **(Figure 1b)** we followed standard PDMS double-casting protocols, which have also been used to produce leaf replicas.^42,45,46,50,56^ However, the high temperatures (∼ 80 °C) required for curing the PDMS to produce the imprint, resulted in the *A. thaliana* leaf samples degrading. Thus, having a dramatic effect on the quality of the imprint. Alternatively, Wu *et al.* imprinted the leaf topography for producing microfluidic channels in PDMS.^43^ Using this protocol the PDMS was prepared at a ratio of 20:1 (base to curing agent), and cured in an oven for 24 h at 45 °C. However, we observed sections of PDMS in contact with the *A. thaliana* leaf samples were not curing. Which we conjecture was due to the moisture content of the leaf samples. As a result, we developed a drying process to minimise the surface moisture content of the leaves. This was undertaken by placing the *A. thaliana* samples affixed to the polystyrene petri dish substrates into either an automatic or a vacuum desiccator. The samples were kept in the desiccator until the surface moisture of the leaves had reduced. This process enabled control over the leaf drying and minimised the degradation of the leaf sample. We also observed that drying the leaf sample prior to affixing the sample to the petri dish resulted in the leaf shrivelling, which rendered the sample unusable. Furthermore, we decreased the curing time to 20 h to minimise degradation of the leaf cuticular waxes. The degradation of the leaf cuticular waxes has been conjectured to result in the leaf remaining attached to the imprint, after peeling the imprint away from the leaf sample.^43^

Any leaf residue remaining on an imprint impacted the quality of the resulting leaf replica. To remove any leaf residue a digestion protocol was utilised. We selected a basic solution of sodium hydroxide and sodium carbonate, as this method was quick in comparison to enzyme digestion protocols. Furthermore, no adverse effects were observed on the PDMS imprints (**Supplementary Information X**).

To produce the PDMS replica leaf surface, the imprint was treated with an anti-adhesion coating. Two different approaches were utilised, where the surface was treated with a hydroxypropylmethylcellulose (HPMC) solution or with trichloro(1H,1H,2H,2H-perfluorooctyl)silane (FDTS).^45,46 50,51^ We found that treatment with FDTS allowed the PDMS leaf replica to be peeled of the leaf imprint easier than those treated with the HPMC solution. We conjecture this is due to a smaller adhesive force of imprints coated with FDTS rather than HPMC.^51^ This is more important for leaf surfaces with an abundance of trichomes, which are inherently fragile and can readily breakoff when the replica leaf is being peeled away from the leaf imprint. However, if one does not have access to a plasma asher/cleaner that is required to produce oxygen plasma for FDTS treatment, then producing a PDMS replica using HPMC as an anti-adhesion coating will suffice. We were unable to observe any apparent differences in replica *A. thaliana* abaxial (lower) surfaces produced by either method (resulted not shown).

Due to the complex nature of trichomes, the fabrication protocol was improved to increase trichome yield. This was achieved by degassing the leaf imprint at various stages of the fabrication protocol for 2 h each time, which assisted in the passive filling of complex microstructures, such as trichomes.^57^ For instance, degassing occurred prior to placing the leaf imprint in the digestion solution. As *A. thaliana*, trichomes are predominantly found on the adaxial (upper) surface of the leaf, these additional steps were only undertaken when replicating the adaxial surface. However, the overall yield for complete trichomes is relatively low, and further improvements need to be made in this area.

The PDMS leaf replica surfaces were produced using PDMS mixed at a ratio of 10:1 (base to curing agent) and cured for 2 h on a hot plate **(Figure 1d)**. Once cooled to room temperature the replica was carefully peeled off the leaf imprint **(Figure 1e)**. The fidelity of the PDMS leaf replicas were examined using optical microscopy and atomic force microscopy (AFM).

### Optical Imaging

To examine the fidelity of our *A. thaliana* PDMS replica leaf surfaces, we initially used optical microscopy (**Figure 2**). The surface topography of leaves vary considerably and different microfeatures can have different structures. Microstructures that can be found on the surface of *A. thaliana* leaves include stomata, trichomes, and grooves.^58,59^ Where, stomata are pores that control transpiration, and enable gas exchange to occur between the atmosphere and the leaf. For instance, stomata enable carbon dioxide to enter the leaf, which is important for photosynthesis to occur.^13,60^ Whereas, trichomes offer different funcationality. Such as minimising water loss from the leaf surface, regulating the temperature of the leaf, reducing the effects of ultraviolet radiation, and/or the secretion of metabolites to deter herbivores and inhibit pathogen development.^13,61^

**Figure 2:**
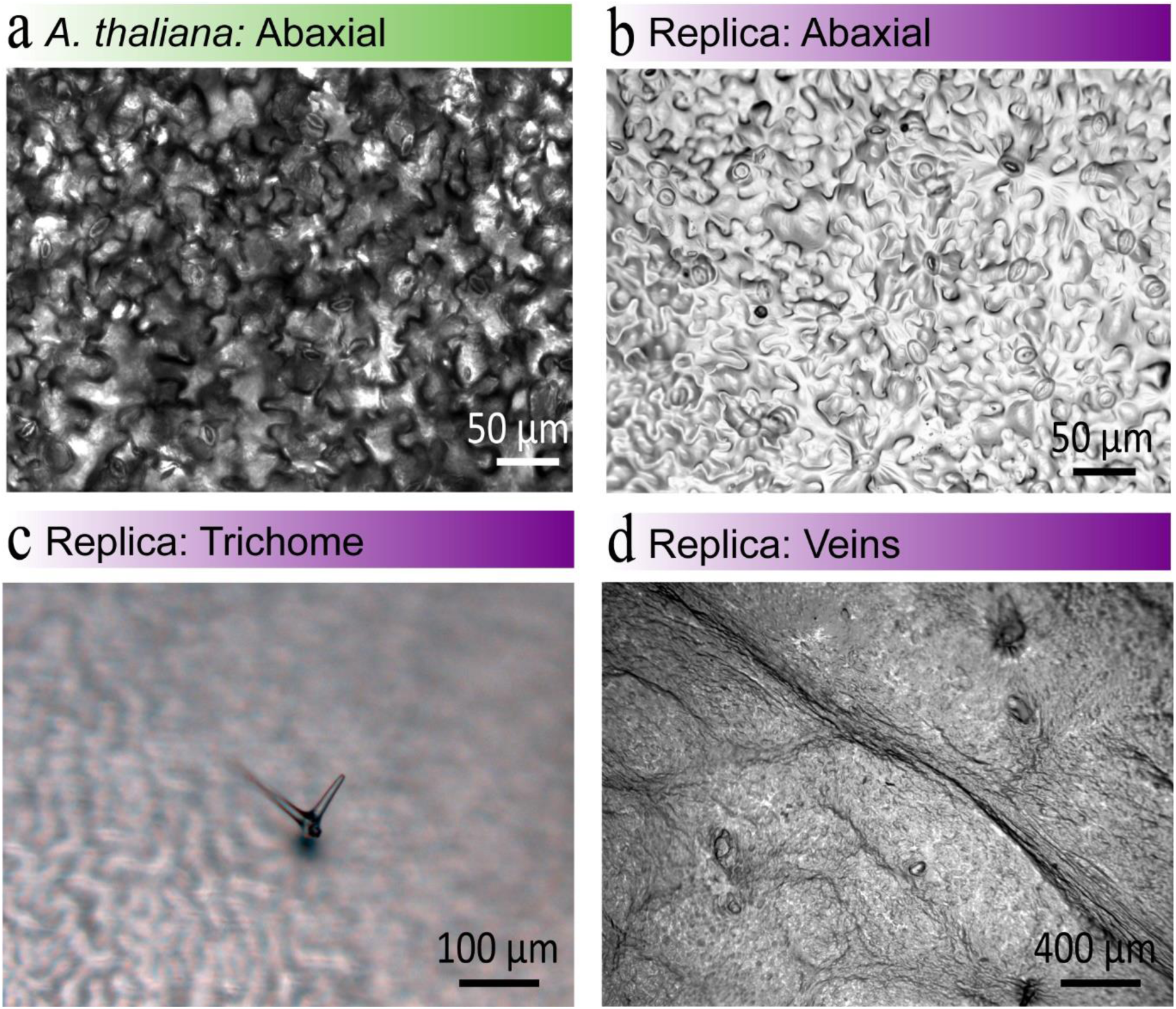
Optical Microscopy Images of *A. thaliana* leaves and replica. (a) A living leaf. (b-d) Replica leaf surfaces, highlighting the different structural aspects of the leaf. Note: abaxial surface is the lower leaf surface, and trichomes are leaf hairs.

For our PDMS replica leaves we were primarily focusing on replicating the abaxial (lower) surface of *A. thaliana* leaves (**Figure 2a-b**). As we were replicating *A. thaliana* leaves for phyllosphere microbiology studies, and bacteria has been found to be more abundant on the abaxial surface, as opposed to the adaxial surface.^37,62,63^ By doing so our work naturally focussed on the fidelity of the replicated stomata and grooves – the most abundant microfeatures on the abaxial surface of *A. thaliana* leaves (**Figure 2a**). The length and width of stomata on the replica *A. thaliana* leaves were measured to be 14.0 ± 1.2 µm and 9.5 ± 1.1 µm, respectively (**Figure 2b**). Which is comparable to the length and width of stomata on living *A. thaliana* leaves, which were measured to be 16.8 ± 2.5 µm and 9.0 ± 1.5 µm, respectively (**Figure 2a**).

A PDMS replica trichome is presented in **Figure 2c**, which highlights the potential of our approach for replicating trichomes found on *A. thaliana* leaves using a double-casting PDMS. Previously the replication of trichomes has only been possible using polyvinylsiloxane imprints in combination with epoxy replicas.^22,24^ A potential future improvement for replicating *A. thaliana* trichomes might be an approach using polyvinylsiloxane imprints, and PDMS replicas. However, negotiating the quick curing time of polyvinylsiloxane would be important for producing a suitable replica.^24,36^ Producing replicas in PDMS is more favourable than using epoxies. This is due to the properties of PDMS, including: biocompatibility, optical transparency, the ease of changing the hydrophobic properties of the surface for attachment studies, and the ability to add fillers for nutrient studies.^64,65^ In addition, the surface energy and bacterial viability on PDMS replica leaf surfaces are comparable to leaf surfaces.^15^

### AFM Imaging

To further investigate the suitability of our replication protocol for producing leaf replicas in PDMS we used atomic force microscopy (AFM) **(Figure 3)**. However, imaging the leaf surfaces with AFM presented some challenges. (1) The roughness of the leaf surface, as an AFM is optimised for measuring the topography of films with nanoscale topography, in addition to nanomechanical and electrochemical characterisation. (2) To obtain high resolution images using AFM can take anywhere from a few minutes to several hours depending on the sample. This was problematic when imaging the *A. thaliana* living leaf samples, as the leaves readily degraded over the course of imaging. Thus, affecting the resolution of the AFM images, and increased the probability of the cantilever tip losing contact. (3) The softness of the *A. thaliana* living leaf, PDMS imprint, and PDMS replica, also influenced the quality of the AFM image. Which resulted in the cantilever not sufficiently tracking the surface of the samples. We minimised this by selecting an appropriate cantilever. (4) In addition, due to the *A. thaliana* living leaf not reflecting the microscope light, this resulted in limited visibility of the leaf surface. Thus, making it difficult to find stomata, and areas with reasonable flatness to allow for imaging.

**Figure 3:**
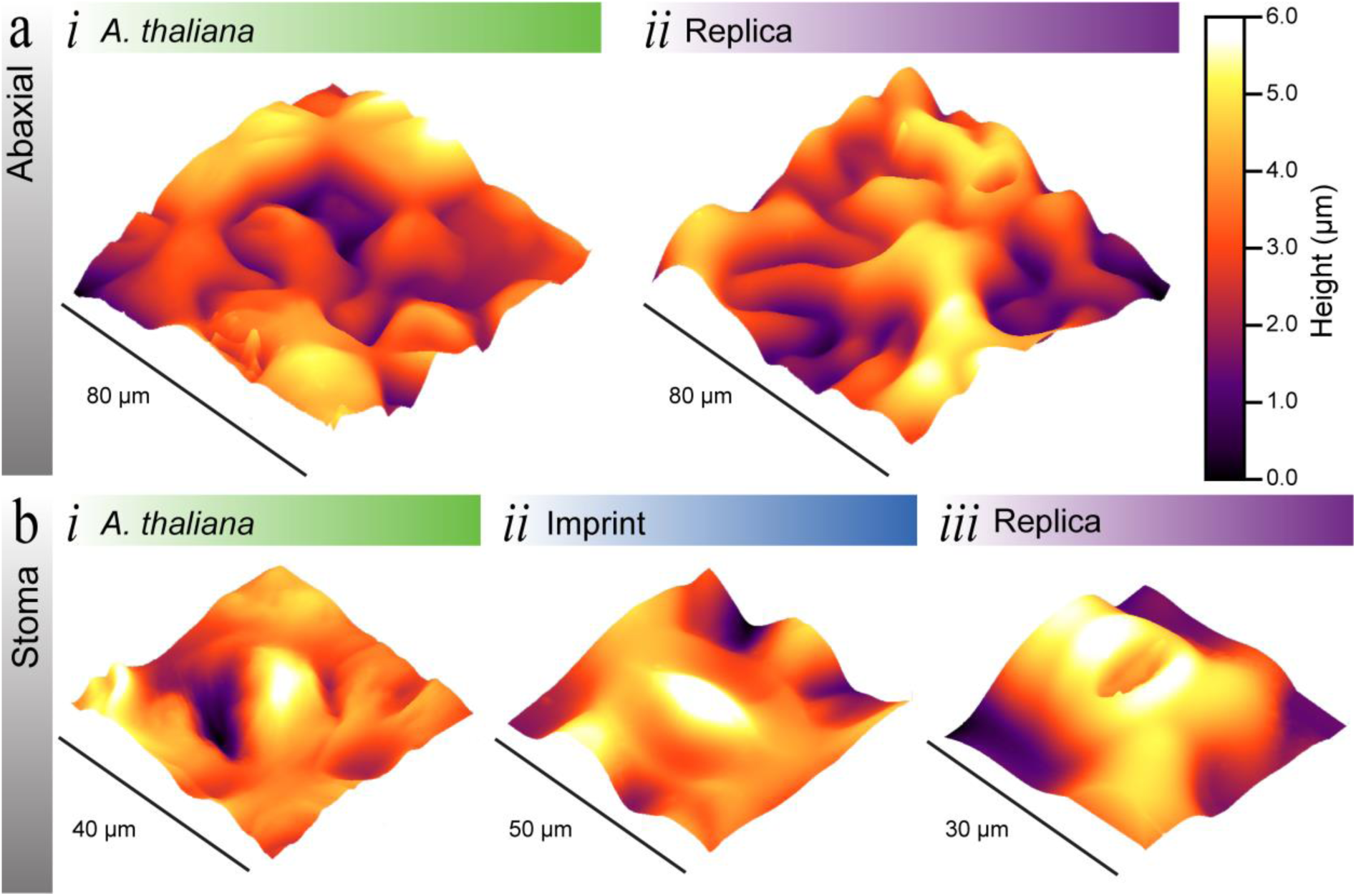
AFM Characterization of A. thaliana Leaf Surfaces. (a) AFM images of the abaxial surface of (i) living and (*ii*) replica A. *thaliana* leaves. (b) AFM images of stoma found on the abaxial surface of (*i*) living, (ii) imprint, and (*iii*) replica A. *thaliana* leaves. A larger area scan of the (*ii*) stoma imprint is displayed in **Supplementary Figure SX.**

To compensate for the roughness of the leaf surfaces, we used an MFP-3D Origin AFM with a large working height range of 15 µm. Furthermore, to minimise the effects of the degradation of the living leaf, the images were undertaken on fresh leaf samples that were gently flattened and affixed to a glass substrate with double-sided tape. In addition, the AFM images were acquired in less than 20 minutes.

As our interest is focused on replicating the abaxial surface, we looked at the fidelity of grooves and stomata, which can be seen in **Figure 3**. The large area topographical AFM images in **Figure 3a** of the *A. thaliana* living leaf and the PDMS replica leaf, display the roughness and variability of the leaf topography. An important microfeature on the abaxial surface of *A. thaliana* are stomata, which can be seen in **Figure 3b**. The images taken of the stoma from a living leaf, leaf imprint, and replica leaf, show similar dimensions. However, like anything in nature the size of stomata varies. In conclusion, the similarities of the microfeatures in the large area topographical AFM images **(Figure 3a)**, and the stomata images **(Figure 3b)** indicate that the PDMS replica *A. thaliana* leaf surface is representative of the *A. thaliana* living leaf surface.

### Contact Angle Measurements

The cuticle, which is a wavy surface, which covers the plant leaf’s epidermis, and protects the plant from the external environment.^66^ Furthermore, the cuticle prevents water, ion, and nutrient loss, whilst preventing pathogenic attacks against the plant host.^67^ An important property of the cuticle is its surface energy, and in particular its hydrophobicity. In phyllosphere microbiology the surface hydrophobicity is important, as the presence of water on the leaf surface influences resources availability, and microorganism colonisation patterns.^68^ In addition, microorganism attachment processes are influenced by the hydrophobicity of the leaf cuticle. Microorganisms can achieve attachment by one of two processes: (1) adapting to enable attachment, or (2) by forming biofilms.^14,69^

The surface energy of a surface can be classified as either hydrophilic, hydrophobic, or superhydrophobic, when the contact angle of water is < 90°, > 90°, and >150°, respectively. The measured contact angle of leaf cuticles/surfaces can vary considerably, from hydrophilic to superhydrophobic.^26,70,71^ We used deionised water to conduct contact angle measurements and recorded droplets with a volume less than 60 µL.

Contact angles were obtained for the adaxial (upper) and abaxial (lower) surfaces of leaf samples from mature *A. thaliana* plants grown in soil **(Figure 3)**. This was undertaken as some plants have differing hydrophobic properties between the adaxial and abaxial surfaces.^70,71^ In the case of *A. thaliana*, no significant difference (N = 5) was observed between the adaxial and abaxial surfaces **(Figure 3)**. A mean contact angle of 97 ±1° for *A. thaliana* was measured; thus, indicating that the surface of *A. thaliana* leaves are hydrophobic.

In addition, contact angle measurements were obtained for *A. thaliana* PDMS replica adaxial and abaxial leaf surfaces **(Figure 3)**. No significant difference (N = 5) was observed between the PDMS replica adaxial and abaxial leaf surfaces. A mean contact angle of 99 ±1° for the PDMS replica leaves was measured; thus, indicating that the surface of the PDMS replica leaves are hydrophobic. Furthermore, no significant difference was observed between the *A. thaliana* and PDMS replicas leaf surfaces.

In addition, PDMS replica surfaces provide the ability to examine the influence of hydrophobicity and the role this has on attachment studies. As the degree of hydrophobicity of a PDMS surface can be temporarily modified with the use of oxygen plasma. This duration can be extended through the use of polyvinylpyrrolidone (PVP) treatment. Oxygen plasma and PVP treatment are not harmful to microorganisms.^65^ Thus, enabling more extensive attachment studies to be undertaken using a PDMS replica leaf surface. This is of importance, as this is a research area that has been highlighted in phyllosphere microbiology that requires more extensive studies to be undertaken.^14^

### Bacterial Visualisation Studies

To examine the suitability of the PDMS replica leaf for phyllosphere microbiology, we used the bacterium *Pantoea agglomerans* 299R as our model microorganism **(Figure 5)**. The bacterium *P. agglomerans* 299 was isolated from a Bartlett pear tree leaf. Strain *P. agglomerans* 299R is a spontaneous rifampicin resistant mutant of *P. agglomerans* 299.^52^ *P. agglomerans* 299R were selected as our model microorganism, as it is: (1) a model microorganism for leaf colonisation studies; (2) well characterised and fully sequenced; and (3) it is genetically amendable (able to produce mutants and bioreporters).^52^ The *P. agglomerans* 299R were modified by genomic Tn7 transposon insertion carrying a mScarlet-1 fluorescent protein. The modified *P. agglomerans* 299R∷MRE-Tn7-145 was selected due to its high brightness.^72^

The bacteria distributions for the living A. *thaliana* **(Figure 5a)** and PDMS replica **(Figure 5b)** abaxial surfaces, were influenced by the distribution of the bacteria suspension droplets. On both *A. thaliana* and the PDMS replica leaf surfaces minimal wetting was observed **(Figure 4)**. Which indicates that the shape of the droplets were predominantly influenced by the leaf microfeatures of the abaxial surface (grooves and stomata). In addition, the bacteria were confined to the droplets as there was no residual surface moisture from the A. *thaliana* or the PDMS replica leaf. Furthermore, more bacteria were observed at the edge of the droplet interface in contact with either the *A. thaliana* or the PDMS replica leaf. Thus, the distribution of bacteria observed on the PDMS replica leaf surface **(Figure 5b)**, was comparable to the distribution observed on the living *A. thaliana* **(Figure 5a)**.

**Figure 4:**
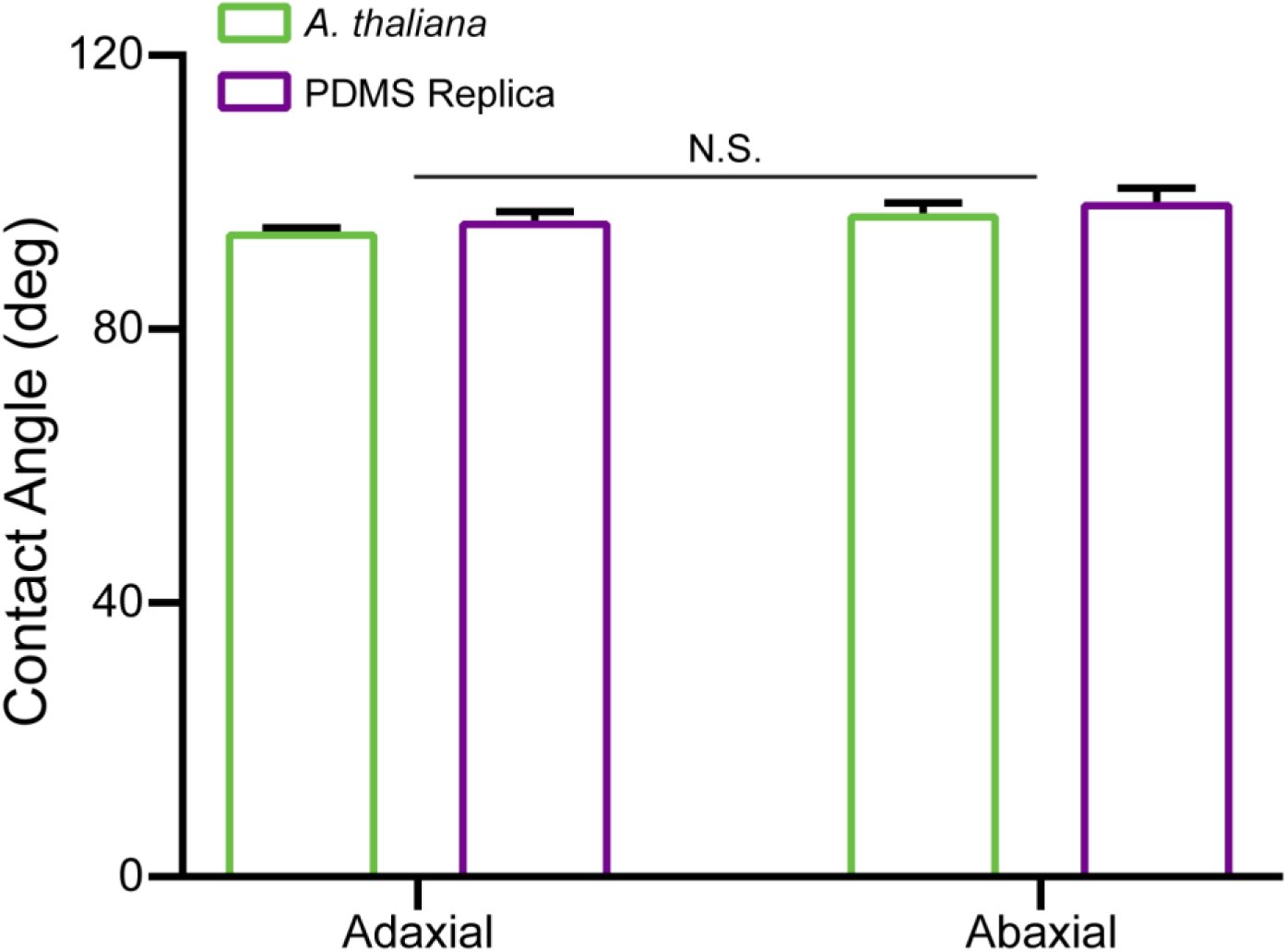
Contact Angle of Living and PDMS Replica A. *thaliana* Leaf Surfaces. Contact angles of living and replica leaf samples, for both the adaxial (upper) and abaxial (lower) surfaces of A. *thaliana*. Results are presented as mean ± SEM (standard error of the mean). N.S. indicates no significant difference between the measurements.

**Figure 5:**
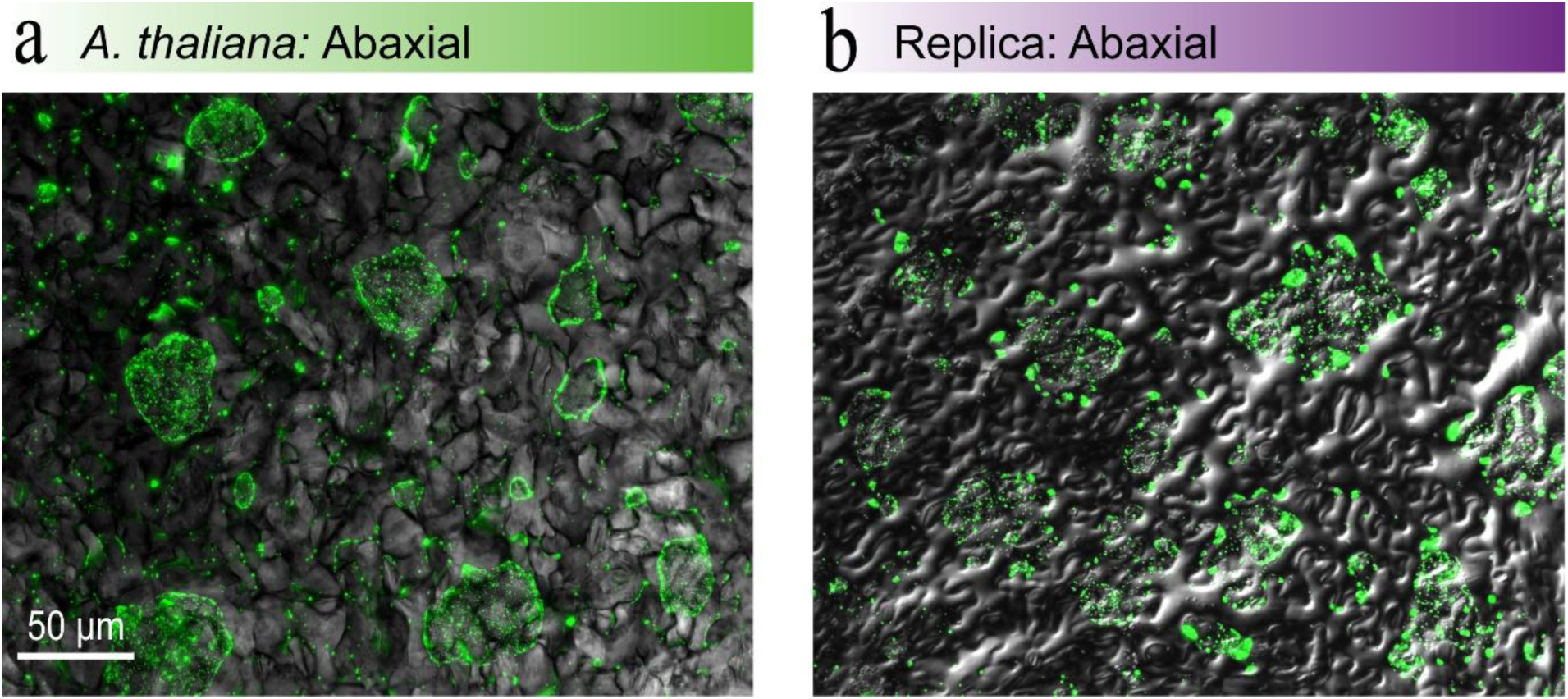
Bacterial Distribution on Abaxial Surfaces. The bacterium *P. agglomerans* 299R∷MRE-Tn7-145 visualised on (a) living *A. thaliana* and (b) PDMS replica abaxial surfaces.

In addition, using the PDMS replica leaf surfaces, results in several advantages over the process used for traditional bacterial visualisation studies undertaken on living leaves. The advantages included: (1) The PDMS replica leaf does not degrade during visualisation studies; (2) The exact same sample can be visualised over several days, which enables time-lapse studies to be undertaken; and (3) as neither mounting resin or cover slides were required, the distribution of the bacteria was not disrupted.

### Conclusions and Future outlook

Our work has demonstrated the potential double-casting PDMS to produce leaf replicas from plants grown under all conditions, including the more challenging optimal case. Based on our observations, other reported replication processes most likely used plants that were grown under non-optimal (stressed) conditions. In contrast, to test the suitability of our replication process we used *A. thaliana* plants grown in optimal conditions in either soil or nutrient agar. This provided a new challenge in form of a reduction of the leaf cuticular waxes and increase in moisture content in the leaves, which influences the curing ability of polymers

In addition, for the first time we have highlighted the potential of replicating the delicate structure of trichomes using PDMS for both the imprint and replica. Previously the replication of trichomes has only been possible using polyvinylsiloxane imprints, in combination with epoxy replicas.

Using microscopy and atomic force microscopy, we have demonstrated that our replication protocol is suitable for replicating the intricate topography of leaf surfaces, for phyllosphere microbiology. Furthermore, the measured surface energy of the living *A. thaliana* leaf surfaces, and the *A. thaliana* PDMS replica surfaces are comparable.

In addition, we examined bacterial distribution using modified *P. agglomerans* 299R∷MRE-Tn7-145 on both living and replica *A. thaliana* abaxial surfaces. The distribution of bacteria observed on the PDMS replica *A. thaliana* abaxial leaf surfaces, were observed to be comparable to the distribution on the living *A. thaliana* abaxial leaf surfaces. Furthermore, using PDMS replica leaf surfaces offer several advantages over living leaves, for example, the PDMS leaf replica surfaces do not degrade, and the exact same sample can be visualised over several days, which enables time-lapse studies of bacteria distributions to be undertaken.

In summary, the results presented here indicate that our replication process for producing replica leaves in PDMS, is suitable for phyllosphere microbiology studies. In our current work, we are investigating nutrient permeability with our PDMS replica *A. thaliana* leaf surfaces, for plant-microbe interactions at a single-cell resolution.

## Acknowledgements

This work was funded by the Biomolecular Interaction Centre and Marsden Grant UOC1704. R.S. thanks the National Science Challenge - Science for Technological Innovation for a Post-Doctoral Fellowship. M.B. is supported by a University of Canterbury Doctoral scholarship. The authors would like to thank Prof. Paula Jameson (The University of Canterbury) for gifting the *Arabidopsis thaliana* seeds, Dr. Jenny Malmstrom and Dr. Yiran An (The University of Auckland) for their assistance with acquiring the AFM scans, and Helen Devereux and Gary Turner for technical assistance.

## References

1 Bauer, S. et al. 25th anniversary article: a soft future: from robots and sensor skin to energy harvesters. Adv. Mater. 26, 149–162 (2014).

2 Chou, H.-H. et al. A chameleon-inspired stretchable electronic skin with interactive colour changing controlled by tactile sensing. Nat. Commun. 6, 8011 (2015).

3 Nakata, T. et al. Aerodynamics of a bio-inspired flexible flapping-wing micro air vehicle. Bioinspir. Biomim. 6, 045002 (2011).

4 Evans, J., Inalhan, G., Jang, J. S., Teo, R. & Tomlin, C. J. in Digital Avionics Systems, 2001. DASC. 20th Conference. 1C3/1-1C3/12 vol. 11 (IEEE).

5 Nishimoto, S. & Bhushan, B. Bioinspired self-cleaning surfaces with superhydrophobicity, superoleophobicity, and superhydrophilicity. RSC Adv. 3, 671–690 (2013).

6 Huh, D., Hamilton, G. A. & Ingber, D. E. From 3D cell culture to organs-on-chips. Trends Cell Biol. 21, 745–754 (2011).

7 Huh, D. et al. Microfabrication of Human Organs-on-Chips. Nat. Protoc. 8, 2135–2157 (2013).

8 Polacheck, W. J., Li, R., Uzel, S. G. & Kamm, R. D. Microfluidic platforms for mechanobiology. Lab Chip 13, 2252–2267, doi:http://dx.doi.org/10.1039/c3lc41393d (2013).

9 Soffe, R. et al. Lateral Trapezoid Microfluidic Platform for Investigating Mechanotransduction of Cells to Spatial Shear Stress Gradients. Sensor. Actuat. A Chem. 251, 963–975 (2017).

10 Nahavandi, S. et al. Microfluidic Platforms for Biomarker Analysis. Lab Chip 14, 1496–1514 (2014).

11 Vorholt, J. A. Microbial Life in the Phyllosphere. Nat. Rev. Microbiol. 10, 828–840 (2012).

12 Meyer, K. M. & Leveau, J. H. Microbiology of the Phyllosphere: A Playground for Testing Ecological Concepts. Oecologia 168, 621–629 (2012).

13 Vacher, C. et al. The Phyllosphere: Microbial Jungle at the Plant–Climate Interface. Annu. Rev. Ecol. Evol. S. 47, 1–24 (2016).

14 Doan, H. K. & Leveau, J. H. Artificial Surfaces in Phyllosphere Microbiology. Phytopathology 105, 1036–1042 (2015).

15 Soffe, R., Altenhuber, N., Bernach, M., Remus-Emsermann, M. N. & Nock, V. Comparison of Replica Leaf Surface Materials for Phyllosphere Microbiology. engrXiv, doi:https://doi.org/10.31224/osf.io/2pzrv.

16 Jacobs, J., Carroll, T. & Sundin, G. The role of Pigmentation, Ultraviolet Radiation Tolerance, and Leaf Colonization Strategies in the Epiphytic Survival of Phyllosphere Bacteria. Microb. Ecol. 49, 104–113 (2005).

17 Rivas, L., Fegan, N. & Dykes, G. A. Attachment of Shiga Toxigenic Escherichia coli to Stainless Steel. Int. J. Food Microbiol. 115, 89–94 (2007).

18 Jeffree, C. E. The Fine Structure of the Plant Cuticle. Annual Plant Reviews: Biology of the Plant Cuticle 23, 11–125 (2006).

19 Melotto, M., Underwood, W., Koczan, J., Nomura, K. & He, S. Y. Plant Stomata Function in Innate Immunity Against Bacterial Invasion. Cell 126, 969–980 (2006).

20 Rusconi, R., Garren, M. & Stocker, R. Microfluidics Expanding the Frontiers of Microbial Ecology. Annu. R. Biophys. 43, 65–91 (2014).

21 Koch, K., Dommisse, A., Barthlott, W. & Gorb, S. N. The use of Plant Waxes as Templates for Micro-and Nanopatterning of Surfaces. Acta Biomater. 3, 905–909 (2007).

22 Koch, K., Schulte, A. J., Fischer, A., Gorb, S. N. & Barthlott, W. A Fast, Precise and Low-Cost Replication Technique for Nano- and High-Aspect-Ratio Structures of Biological and Artificial Surfaces. Bioinspir. Biomim. 3, 046002(046001–046010) (2008).

23 Zhang, B. et al. Fabrication of Biomimetically Patterned Surfaces and their Application to Probing Plant–Bacteria Interactions. ACS Appl. Mater. Inter. 6, 12467–12478 (2014).

24 Schulte, A. J., Koch, K., Spaeth, M. & Barthlott, W. Biomimetic replicas: transfer of complex architectures with different optical properties from plant surfaces onto technical materials. Acta Biomater. 5, 1848–1854 (2009).

25 Lepore, E. & Pugno, N. Superhydrophobic Polystyrene by Direct Copy of a Lotus Leaf. BioNanoScience 1, 136–143 (2011).

26 Sun, M. et al. Artificial Lotus leaf by Nanocasting. Langmuir 21, 8978–8981 (2005).

27 Wang, B., Liang, W., Guo, Z. & Liu, W. Biomimetic super-lyophobic and super-lyophilic materials applied for oil/water separation: a new strategy beyond nature. Chemical Society Reviews 44, 336–361 (2015).

28 Heaton, J. C. & Jones, K. Microbial Contamination of Fruit and Vegetables and the Behaviour of Enteropathogens in the Phyllosphere: A Review. J. Appl. Microbiol. 104, 613–626 (2008).

29 Compant, S., Van Der Heijden, M. G. & Sessitsch, A. Climate Change Effects on Beneficial Plant–Microorganism Interactions. FEMS Microbiol. Ecol. 73, 197–214 (2010).

30 Newton, A., Gravouil, C. & Fountaine, J. Managing the Ecology of Foliar Pathogens: Ecological Tolerance in Crops. Ann. Appl. Biol. 157, 343–359 (2010).

31 Herman, K., Hall, A. & Gould, L. Outbreaks Attributed to Fresh Leafy Vegetables, United States, 1973–2012. Epidemiol. Infect. 143, 3011–3021 (2015).

32 Painter, J. A. et al. Attribution of Foodborne Illnesses, Hospitalizations, and Deaths to Food Commodities by using Outbreak Data, United States, 1998–2008. Emerg. Infect. Dis. 19, 407–415 (2013).

33 Doyle, M. & Erickson, M. Summer Meeting 2007 – The Problems with Fresh Produce: An Overview. J. Appl. Microbiol. 105, 317–330 (2008).

34 Olaimat, A. N. & Holley, R. A. Factors Influencing the Microbial Safety of Fresh Produce: A Review. Food Microbiol. 32, 1–19 (2012).

35 Althaus, D., Hofer, E., Corti, S., Julmi, A. & Stephan, R. Bacteriological Survey of Ready-to-Eat Lettuce, Fresh-Cut Fruit, and Sprouts Collected from the Swiss Market. J. Food Protect. 75, 1338–1341 (2012).

36 Prüm, B., Bohn, H. F., Seidel, R., Rubach, S. & Speck, T. Plant surfaces with cuticular folds and their replicas: influence of microstructuring and surface chemistry on the attachment of a leaf beetle. Acta Biomater. 9, 6360–6368 (2013).

37 Remus-Emsermann, M. N. et al. Spatial distribution analyses of natural phyllosphere-colonizing bacteria on Arabidopsis thaliana revealed by fluorescence in situ hybridization. Environ. Microbiol. 16, 2329–2340 (2014).

38 Koornneef, M. & Meinke, D. The development of Arabidopsis as a model plant. The Plant Journal 61, 909–921 (2010).

39 Poorter, H. et al. The art of growing plants for experimental purposes: a practical guide for the plant biologist. Functional Plant Biology 39, 821–838 (2012).

40 Cameron, K. D., Teece, M. A. & Smart, L. B. Increased accumulation of cuticular wax and expression of lipid transfer protein in response to periodic drying events in leaves of tree tobacco. Plant physiology 140, 176–183 (2006).

41 Park, S., van Rijn, P. & Böker, A. Artificial leaves via reproduction of hierarchical structures by a fast molding and curing process. Macromolecular rapid communications 33, 1300–1303 (2012).

42 McDonald, B., Patel, P. & Zhao, B. Micro-structured Polymer Film Mimicking the Trembling Aspen Leaf. Chem. Eng. Process Tech. 1, 1012(1011–1018) (2013).

43 Wu, W., Guijt, R. M., Silina, Y. E., Koch, M. & Manz, A. Plant leaves as templates for soft lithography. RSC Adv. 6, 22469–22475 (2016).

44 Mutreja, I. et al. Positive and Negative Bioimprinted Polymeric Substrates: New Platforms for Cell Culture. Biofabrication 7, 025002(025001–025013) (2015).

45 Yang, L., Hao, X., Wang, C., Zhang, B. & Wang, W. Rapid and Low Cost Replication of Complex Microfluidic Structures with PDMS Double Casting Technology. Microsyst. Technol. 20, 1933–1940 (2014).

46 Gitlin, L., Schulze, P. & Belder, D. Rapid Replication of Master Structures by Double Casting with PDMS. Lab Chip 9, 3000–3002, doi:10.1039/B904684D (2009).

47 Podar, D. in Plant Mineral Nutrients 23–45 (Springer, 2013).

48 Babu, A., Kumaresan, G., Raj, V. A. A. & Velraj, R. Review of leaf drying: Mechanism and influencing parameters, drying methods, nutrient preservation, and mathematical models. Renewable and Sustainable Energy Reviews 90, 536–556 (2018).

49 He, J. et al. Fabrication of Nature-Inspired Microfluidic Network for Perfusable Tissue Constructs. Advanced healthcare materials 2, 1108–1113 (2013).

50 Zhuang, G. & Kutter, J. P. Anti-stiction coating of PDMS moulds for rapid microchannel fabrication by double replica moulding. Journal of Micromechanics and Microengineering 21, 105020 (2011).

51 Bhushan, B., Hansford, D. & Lee, K. K. Surface modification of silicon and polydimethylsiloxane surfaces with vapor-phase-deposited ultrathin fluorosilane films for biomedical nanodevices. Journal of Vacuum Science & Technology A: Vacuum, Surfaces, and Films 24, 1197–1202 (2006).

52 Remus-Emsermann, M. N., Kim, E. B., Marco, M. L., Tecon, R. & Leveau, J. H. Draft Genome Sequence of the Phyllosphere Model Bacterium Pantoea agglomerans 299R. Genome Announc. 1, e00036–00013 (2013).

53 Remus-Emsermann, M. N., Tecon, R., Kowalchuk, G. A. & Leveau, J. H. Variation in Local Carrying Capacity and the Individual Fate of Bacterial Colonizers in the Phyllosphere. ISME J. 6, 756–765 (2012).

54 Forster, B., Van De Ville, D., Berent, J., Sage, D. & Unser, M. Complex wavelets for extended depth-of-field: A new method for the fusion of multichannel microscopy images. Microscopy research and technique 65, 33–42 (2004).

55 Innerebner, G., Knief, C. & Vorholt, J. A. Protection of Arabidopsis thaliana Against Leaf-Pathogenic Pseudomonas syringae by Sphingomonas Strains in a Controlled Model System. Appl. Environ. Microb. 77, 3202–3210 (2011).

56 McDonald, B., Shahsavan, H. & Zhao, B. Biomimetic Micro-Patterning of Epoxy Coatings for Enhanced Surface Hydrophobicity and Low Friction. Macromolecular Materials and Engineering 299, 237–247 (2014).

57 Hosokawa, K., Sato, K., Ichikawa, N. & Maeda, M. Power-free poly (dimethylsiloxane) microfluidic devices for gold nanoparticle-based DNA analysis. Lab Chip 4, 181–185 (2004).

58 Eckerson, S. H. The Number and Size of the Stomata. Bot. Gaz. 46, 221–224 (1908).

59 Huchelmann, A., Boutry, M. & Hachez, C. Plant glandular trichomes: natural cell factories of high biotechnological interest. Plant physiology 175, 6–22 (2017).

60 Koman, V. B. et al. Persistent drought monitoring using a microfluidic-printed electro-mechanical sensor of stomata in planta. Lab Chip 17, 4015–4024 (2017).

61 Beck, C. B. An introduction to plant structure and development: plant anatomy for the twenty-first century. (Cambridge University Press, 2010).

62 Krimm, U., Abanda-Nkpwatt, D., Schwab, W. & Schreiber, L. Epiphytic microorganisms on strawberry plants (Fragaria ananassa cv. Elsanta): identification of bacterial isolates and analysis of their interaction with leaf surfaces. FEMS Microbiol. Ecol. 53, 483–492 (2005).

63 Peredo, E. L. & Simmons, S. L. Leaf-FISH: microscale imaging of bacterial taxa on phyllosphere. Front. Microbiol. 8, 2669 (2018).

64 Xue, C. et al. A Carbon Nanotube Filled Polydimethylsiloxane Hybrid Membrane for Enhanced Butanol Recovery. Sci. Rep. 4, 5925(5921–5927) (2014).

65 Hemmilä, S., Cauich-Rodríguez, J. V., Kreutzer, J. & Kallio, P. Rapid, Simple, and Cost-Effective Treatments to Achieve Long-Term Hydrophilic PDMS Surfaces. Appl. Surf. Sci. 258, 9864–9875 (2012).

66 Schreiber, L. Transport Barriers made of Cutin, Suberin and Associated Waxes. Trends Plant Sci. 15, 546–553 (2010).

67 Aragón, W., Reina-Pinto, J. J. & Serrano, M. The Intimate Talk Between Plants and Microorganisms at the Leaf Surface. J. Exp. Bot. 68, 5339–5350 (2017).

68 Leveau, J. H. & Lindow, S. E. Appetite of an epiphyte: quantitative monitoring of bacterial sugar consumption in the phyllosphere. Proceedings of the National Academy of Sciences 98, 3446–3453 (2001).

69 Morris, C. E. & Monier, J.-M. The Ecological Significance of Biofilm Formation by Plant-Associated Bacteria. Annu. Rev. Phytopathol. 41, 429–453 (2003).

70 Brewer, C., Smith, W. & Vogelmann, T. Functional Interaction between Leaf Trichomes, Leaf Wettability and the Optical Properties of water Droplets. Plant Cell Environ. 14, 955–962 (1991).

71 Wang, H., Shi, H., Li, Y. & Wang, Y. The Effects of Leaf Roughness, Surface Free Energy and Work of Adhesion on Leaf Water Drop Adhesion. Plos One 9, e107062(107061–107010) (2014).

72 Schlechter, R. O. et al. Chromatic Bacteria - A broad host-range plasmid and chromosomal insertion toolbox for fluorescent protein expression in bacteria. Under Consideration.

